# Sub-lethal pesticide exposure facilitates the potential northward range shifts of ticks by increasing cold tolerance and overwintering survival

**DOI:** 10.1101/2025.04.14.648794

**Authors:** Kennan J. Oyen, Thomas Arya, Benjamin Davies, Elise M. Didion, Joshua B. Benoit

## Abstract

Pesticides are a primary tool for the control of harmful insects or other organisms throughout the world. Although most organisms are likely to encounter pesticides in the environment, little is known about the effects of these chemicals on the physiology of off-target species. We measured the impact of sublethal pesticide exposure on the cold tolerance of two common tick species. We predicted that sublethal pesticide exposure would make ticks more sensitive to extreme temperatures. In contrast, we found that exposure to sublethal pesticides had the opposite effect, increasing survival at low temperatures. Sublethal pesticide exposure decreased the LT50 (lethal temperature at which 50% mortality is observed) of adult and nymphal *D. variabilis* from −16°C to −19°C and improved overwintering survival. Evaluation of previously available RNA-seq data between rapid cold hardening and pesticide exposure indicated potential transcriptional shifts associated with cross-tolerance between cold and pesticide exposure. To investigate the population-level impacts of this physiological shift in cold tolerance, we developed a novel approach for incorporating physiological survival data into spatially explicit species distribution models (SDMs). Using this approach, we demonstrate that pesticide-induced increases in cold tolerance may permit faster northward range shifts of *D. variabilis*. We also demonstrate that incorporating physiological data into SDMs moderates the estimated impacts of environmental change and may provide more accurate predictions of species responses to changing environments. Although our study is limited to ticks, other studies have shown the effects of pesticides on thermal tolerance traits in diverse species. The extensive use of pesticides may drive complex interactions between species and their environments, leading to altered thermal tolerance traits and establishment in new habitats.

## INTRODUCTION

As the environment changes, the earth is experiencing a global reorganization of species, including geographic range shifts, contractions, and expansions (Eriksson & Hillebrand, 2019). Species distribution models (SDMs) have provided a valuable tool for predicting the impact of environmental change on organisms and a significant body of work incorporates emissions scenarios and occurrence data to predict the future distributions of species. Although useful for hypothesis testing, these purely correlative SDMs may prove spatially and temporarily inaccurate - underestimating the rate of an invasion into new habitats or overestimating a species’ available niche (A. Lee-Yaw et al., 2022; Barbet-Massin et al., 2018; Beale & Lennon, 2012). The reason for the mismatch between predicted responses to environmental change and actual species distributions is likely multifaceted, but one explanation is that temperature has non-linear effects on organism physiology (Buckley & Huey, 2016). These non-linear effects include the additive, synergistic, or antagonistic impacts of multiple abiotic and biotic stressors, which organisms experience in the environment but are rarely included in models that estimate the impacts of temperature variation (Kaunisto et al., 2016).

Abiotic and biotic stressors may include habitat loss, extreme climatic events, infection, or chemical exposure (Coors & De Meester, 2008). These stressors are rarely experienced independently and instead overlap temporally and spatially. Pests are often the target of chemical control measures and exposure to toxic chemicals likely occurs simultaneously with exposure to extreme temperatures. For some arthropods, the impact of chemical exposure in combination with thermal stress results in unpredictable outcomes. For example, in bumblebees, sub-lethal exposure to glyphosate, an organophosphorus herbicide, decreases thermogenesis and leads to the inability to thermoregulate colonies, which is critical for growth and development of eggs and larval stages (Weidenmüller et al., 2022). True bugs (*Apolygus lucorum*), produce heat shock protein (HSP) 90 in response to both high temperatures and pesticides, resulting in greater survival at physiologically hot temperatures in pesticide-exposed individuals (Sun et al., 2014). Several possible mechanisms may underlie these responses. At the cellular level, physiological stressors such as infection, chemical exposure, and thermal shock may have overlapping effects including shared signaling pathways (cross-talk) or shared protective mechanisms (cross-tolerance) (El-Saadi et al., 2023).

Cross-talk and cross-tolerance in arthropods are mechanisms that allow organisms to respond to multiple environmental stressors simultaneously (Boardman, 2024). For instance, temperate and polar insects experience both cold and desiccation stress during winter, and the cellular mechanisms that protect against cold stress also confer protection against desiccation, illustrating cross-tolerance (Sinclair et al., 2013). In contrast, cross-talk is observed when low-temperature stress leads to the upregulation of immune responses, even though there is little mechanistic overlap between cold stress and immune stress at the cellular level (Ferguson et al., 2016). Key signaling pathways involved in cross-talk in arthropods include the immune deficiency (Imd), Toll, and Janus kinase/signal transducers and activators of transcription (JAK-STAT) pathways. These pathways are crucial for the insect immune response, particularly in defense against pathogens (El-Saadi et al., 2023). Cross-talk between these pathways allows arthropods to mount a coordinated response to multiple threats, and likely has broad implications for vectors that can carry and transmit numerous pathogens.

Ticks are among the most prolific vectors of both bacterial and viral pathogens because they are broadly distributed and generalist blood feeders. Recent studies have shown that ticks are expanding their already extensive geographic ranges and exploiting new host populations, a trend likely facilitated by environmental change and increased host movement across the globe (Couper et al., 2021; Porretta et al., 2013). However, the rapid invasion of ticks into new habitats has substantially outpaced some climate model predictions (Estrada-Peña, 2002). The reason for this mismatch between predicted and current range limits of ticks is probably multifaceted, but could be driven by interactions between stressors. Temperature-induced expression of antimicrobial peptides (AMPs) has been shown in *Drosophila,* which may also increase resistance to infection (Štětina et al., 2019). The production of AMPs is initiated by the Imd and Toll pathways which defend against bacterial infection. Conversely, *Borrelia bordgorferi*, the causative agent of Lyme disease, stimulates upregulation of antifreeze glycoproteins which enhances cold tolerance and may explain increased overwintering survival in northern climates (Neelakanta et al., 2010). Recent data have shown that infected ticks are twice as likely to survive overwintering compared with uninfected ticks (Nabbout et al., 2023). This highlights the importance of understanding the interactions between multiple stressors, particularly for disease vectors that are often infected with pathogens, exposed to extreme temperatures, and targeted by pesticides.

Resistance to pesticides and thermal stress in arthropods is likely interconnected through mechanisms of cross-tolerance and cross-protection, where exposure to one stressor influences the response to another. Studies have shown that insects, such as mosquitoes and beetles, can develop cross-tolerance between heat and insecticides. For instance, when mosquitoes are exposed to heat shock, they exhibit increased resistance to insecticides like propoxur and permethrin (Mack & Attardo, 2024). Similarly, the exposure of insects to elevated temperatures can enhance their survival against subsequent insecticide exposure, as observed in pests like the diamondback moth and wheat aphid (Perrin et al., 2022; Xing et al., 2023). These interactions are influenced by HSPs, which play a role in both thermal tolerance and insecticide resistance, although the exact mechanisms remain to be fully elucidated. Understanding these dynamics is crucial in the context of environmental change, as changing temperatures may alter the efficacy of insecticides and impact pest management strategies and future distributions of ticks and associated pathogens.

We tested the effect of two pesticides on tick cold tolerance by exposing two species of ticks to sublethal doses of pesticides and measured their cold tolerance. We chose two pesticides which are commonly used throughout residential and agricultural settings: propoxur and chlorpyrifos. Both pesticides act on highly conserved ionoregulatory channels that control the CNS which undergoes systemic depolarization during cold exposure (Robertson et al., 2017). We found that there is a synergistic effect of pesticide and low temperatures whereby sublethal doses of pesticides increase tick cold tolerance across short (days to weeks) and long (overwintering) timeframes. Re-evaluation of previous RNA-seq data sets highlights overlapping transcriptional responses between cold and pesticide exposure. Given the overall increase in general pesticide use throughout the world, this effect could explain the relatively rapid northward invasion of ticks beyond what is predicted by models alone. To test how pesticide-induced increases in cold tolerance may impact tick distributions, we developed physiological species distribution models (pSDMs) by incorporating thermal performance curves as Bayesian priors to correlative SDMs using generalized linear models (GLMs). Incorporating physiological data, significantly improved model fitting. We suggest that the influence of multiple stressors, which can have additive, synergistic, or antagonistic effects, are a potential caveat to current models estimating future changes in tick distributions. Our work highlights the importance of measuring multiple stressors and using physiological data to inform predictive models.

## METHODS

### Study animals

Unfed ticks from each life stage of both *Rhipicephalus sanguineus* and *Dermacentor variabilis* were obtained from Ecto Services Inc. (Henderson NC, USA) as eggs, engorged larvae, and nymphs. After hatching or molting, larvae, nymphs, and adults were used within 2-3 months to reduce any impacts associated with starvation or age (Rosendale et al., 2019). Ticks were maintained at 93% relative humidity (RH), 22 ± 1°C, and 12:12 hr, light:dark (L:D).

### Pesticide application

Chlorpyrifos methyl and propoxur were obtained from Chem Services Inc. with 99.7% and 99.9% purity respectively. Both were dissolved in a 100% acetone to obtain stock solutions of 1.0 mg mL−1 and kept at −18 °C. Experimental solutions were then prepared by serial dilution with ultrapure deionized water to 2.5, 5.0, 10.0, 15.0, and 20.0 PPM based on previous studies (Pathak et al., 2022). Both pesticides were applied using either the larval packet test (Stone and Haydock 1962) or submerging adults in 1.5mL microcentrifuge tubes with pesticide solutions for 20 seconds (Drummond et al., 1973). For the larval packet test, larvae were placed into envelopes folded from filter paper to measure 6 x 6cm and sealed with paper clips. Then 90 ul of treatment or control solutions were pipetted onto the envelope. Ticks were treated with the solvent (acetone diluted to 2 PPM with deionized water) in control groups and 2 PPM of chlorpyrifos or propoxur in treatment groups.

### Cold tolerance testing

To determine the impact of pesticide exposure on cold tolerance, survival to cold exposure was determined using a 2-hour exposure to specific temperatures ranging from −28°C to −16°C (Rosendale et al., 2016). Groups of larvae or individual nymphs, and adults were placed in 1.5cm pcr tubes with mesh lids and then placed in 50ml conical tubes, and suspended in a programmable, refrigerated bath (Arctic A25; Thermo Scientific, Pittsburgh, PA, USA) which maintained the temperature setpoint (±0.1 °C). Following a 10-minute hold, which allowed the conical tube temperature to equilibrate to the set temperature, the bath was held for 2 hours. All ticks were assessed for survival at 24 hours post cold exposure and scored on a scale of 1-4 with the following determinations:

1. No visible effect of cold exposure
2. Slightly impaired; legs somewhat curled but able to move towards and grasp experimenter
3. Alive but heavily impaired. Unable to move towards or grasp experimenter
4. Unmoving and legs adducted

For this study, ticks assessed as 3 or 4 are considered dead and those scored as 1 and 2 are considered alive.

### Overwintering survival

To measure the impact of pesticide exposure on overwintering survival, we compared two geographic locations that are characteristic of northern (Crooks, SD, USA) and southern (Cincinnati, OH, USA) overwintering conditions within the geographic range for *D. variabilis*. 200 adult *D. variabilis* were placed in exclosures in Crooks, SD, USA (North) and Cincinnati, OH, USA (South) from September 2020 through April 2021. Adult *D. variabilis* (even numbers of males and females) were dosed with 20 PPM of chlorpyrifos, propoxur, or solvent and placed in conical tubes with holes drilled in the lid. Tubes were partially filled with soil from each respective field site for substrate and then placed into 1 L aluminum coffee cans filled with local soil. Two HOBO U23 Pro V2 Temperature/Relative Humidity Data Loggers (Onset Computer Corporation, Bourne, MA, USA) were placed in each coffee can with the conical tubes and recorded temperature and relative humidity at 15-minute intervals (data available in Figure S1). Each coffee can with 6 groups of 100 (50:50 male: female) was wrapped in chicken wire to prevent animal predation or interference, and buried to rim height. Once buried, cans were left undisturbed for approximately 8 months. After this period, the cans were removed and the number of ticks alive in each vial was assessed.

### Comparative RNA-seq studies between cold and pesticide exposure

To determine potential mechanisms for the overlap between cold and pesticide exposure, we compared the results of two previous studies that focused on these topics, which examined the transcriptional response to pesticides and rapid cold hardening (Pathak et al., 2022; Rosendale et al., 2022). Reads were mapped to the *de novo* assembly (Rosendale et al. 2022) using CLC Genomics with at least 70% of the read having at least 80% identity with the reference and a mismatch cost of 2. Expression values were measured as transcript per million mapped reads (TPM). Statistical analyses were performed using an EDGE test to identify the transcripts that were significantly enriched using a false discovery rate (FDR) adjusted p-value cut-off of < 0.05. To identify key tissue-specific biological processes, GO analysis was performed on g:Profiler using *Dermacentor andersoni* as a proxy for the corresponding *D*. *variabilis* contigs in each transcriptome (Kolberg et al., 2023). Treemaps were constructed for both tissues based on enriched GO categories with Revigo (Supek et al., 2011). Illumina sequencing datasets are available in association with the following NCBI Birojects - PRJNA657863 and PRJNA783667.

### Modeling

#### Climatic Data

The 19 “Bioclim” variables represent the average conditions of temperature and rainfall for the years 1970–2000 (Hijmans et al., 2005). The variables used in this study correspond to the standard 19 “Bioclim” variables which were extracted from the WorldClim database using the sdmpredictors package in R (http://www.worldclim.com/). We selected climate variables that are biologically relevant to *D. variabilis* including: Elevation, BIO 2 (mean diurnal range), BIO 3 (isothermality), BIO 5 (max temperature of warmest month, BIO 6 (min temperature of coldest month), BIO 13 (precipitation of wettest month), and BIO 15 (precipitation seasonality) (James et al., 2015; Minigan et al., 2018).

#### Occurrence data

In total, models were fitted to 425,390 occurrence data points for *D. variabilis* from the Global Biodiversity Database (GBIF). Data was cleaned using the scrubr package which eliminated occurrences with reported uncertainty greater than a 10×10 km^2^ area. Environmental predictors were obtained using the sdmpredictors package from WorldClim. We started with the first 20 layers and masked and cropped those layers to the modeling region of interest, which contained all occurrence records for established *D. variabilis* populations, excluding sites throughout Alaska, as those are likely point occurrences rather than established populations (Hahn et al., 2020). We built a data frame of all the cleaned species occurrence data, gridded to a resolution of 2.5 arcminutes (∼ 5km), with one row per pixel with the value of the variables and the presence/absence of species records. Therefore, each presence or absence was associated with a specific set of environmental variables.

#### Physiological Sub Model

To incorporate the temperature responses, we fit a logistic curve to the survival data with the slope and intercept from a logistic model to create a physiological sub-model (Figure S2a,b). The logistic regression model has the form:

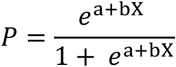

where P is the probability of an event (e.g., the probability of survival) and e is the basis of the natural logarithm, and a and b are the parameters of the model (in this case the slope and regression of the best fit line for the survival data). This resulted in two separate survival functions for pesticide-exposed ticks and unexposed ticks. For heat tolerance, we modeled data from Fieler et al, which tested *D. variabilis* thermal tolerance from 35°C to 55°C (Fieler et al., 2021). Using this data, we established the upper lethal temperature at which 50% mortality occurs (ULT50) to be 39.65°C. This sigmoidal curve was then applied to every pixel of the max temperature of the warmest month (BIO5) and min temperature of the coldest month (BIO6) which yielded a map of survivorship as a function of temperature for each pixel. For instance, a pixel of BIO5 at a temperature of 30.7 was transformed to a value of 0.54 in the new raster layer corresponding to approximately 54% survival at this temperature (Rodríguez et al., 2019). These values were then used to create a physiological species distribution model (pSDM). To construct the pSDM, species occurrences were fitted to environmental predictors using Bayesian GLMs. The posterior distribution of coefficients from the physiological sub-model were used as priors for each of the GLMs. Species occurrences were treated as Bernoulli trials, where p is the probability of finding the species (relative suitability).

### Analysis

The proportion of larvae that survived in each tube was recorded with the mean ± SEM reported for each treatment in the figures. The survival data were analyzed within each species and life stage using generalized linear mixed effects models from the lme4 package with the proportion survived in each tube as the response variable, the treatment as the fixed effect and trial number as a random intercept. Post hoc comparisons of significant models were conducted using the emmeans package in r with the lmerTest package for pairwise comparisons and degrees of freedom were calculated using the Satterthwaite method. Significance was judged at P < 0.05. The temperature resulting in 50% mortality was calculated using a dose.p function in the MASS package. All tests were made using the R^©^ Software v. 3.6.1 (R Core Team, 2017).

For both the SDMs and pSDMs we estimated cross-validation model performance from the training data to the testing data using the Boyce index (Hirzel et al., 2006). The Boyce index ranges from −1 to 1, where 0 means the model does not differ from random and 1 indicates a perfect fit to the data (Hirzel et al. 2006). We assessed model performance using multiple metrics including 1) the Area Under the precision recall curve (AUC); 2) Area under the receiver operating characteristic curve (AUC_ROC_); 3) the True Skill Statistic (TSS); 4) Miller’s calibration slope; 5) CCR (correct classification rate); and, 6) measures of sensitivity, specificity, precision, and recall, using the optim.thresh function in the SDMTools package in R (VanDerWal et al., 2014). We quantified the degree to which species distribution changes between: 1) current and future climate, and 2) regular SDMs and pSDMs. These changes were quantified using the Schoener’s D and Hellinger’s I matrices of overlap (Warren et al., 2008). The D and I matrices measure the overall match between the species distributions and range from 0, no overlap, to 1, identical distributions (Broennimann et al., 2012; Warren et al., 2010).

## RESULTS

### Pesticides increase survival at low temperatures

Exposure to pesticides increased survival at −16°C in both adult *D. variabilis* (Figure 1a; chlorpyrifos: estimate = −0.1923, *p*=0.053; propoxur: estimate = 0.2048, *p*=0.040) and *R. sanguineus* ticks (Figure 1b; chlorpyrifos: estimate = −0.0225, *p*=0.118; propoxur: estimate = 0.450, *p*=0.007). *Dermacentor* larvae also had higher survival in groups treated with pesticides prior to cold exposure (Figure 1c; chlorpyrifos: estimate = −0.51, *p*=0.0065; propoxur: estimate = −0.37, *p*=0.0337). The same pattern was present for *R. sanguineus* larvae with both propoxur and chlorpyrifos treated groups having higher survival than solvent treated counterparts (Figure 1d; chlorpyrifos: estimate = −0.58, *p*=0.0004; propoxur: estimate = −0.37, *p*=0.0074). Across all larvae and adults, there was no significant difference in survival between propoxur and chlorpyrifos treated groups except in adult *R. sanguineus* which had significantly higher survival in the propoxur treated group compared with the solvent and chlorpyrifos group, which were not significantly different (Fig 1B; chlorpyrifos – solvent: estimate = −0.225, *p*=0.1180). There was no significant difference among ticks treated with solvents, water, or untreated ticks (Figure S3; water - untreated: estimate = −0.0312, *p*=0.8454; solvent - untreated: estimate = −0.0101, *p*=0.9804; water – solvent: estimate = 0.0211, *p*=0.9175).

**Figure 1:**
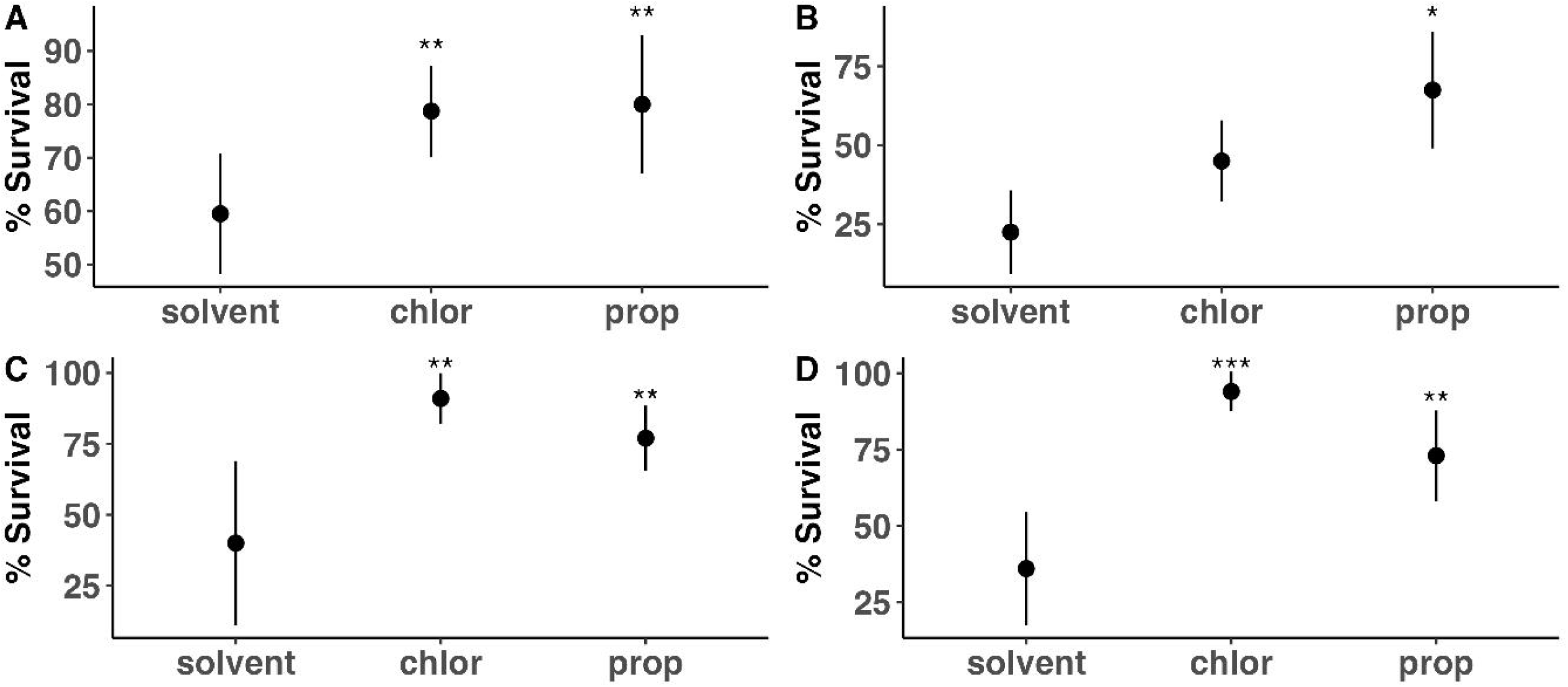
The proportion of adult *Dermacentor* (A) and *Rhipicephalus* (B) and larval *Dermacentor* (C) and *Rhipicephalus* (D) that survived a 2-hour exposure to −16°C following treatment with chlorpyrifos (chlor), propoxur (prop), or a solvent. See text for additional statistical details. Data points represent the mean survival of 5 repeated trials with 20 individuals per trail for adults and 100 for larvae with standard error. Asterisks indicate values that were significantly different from solvent controls (t-test, Satterthwaite’s method, \**P* < 0.05, \*\**P* < 0.01, \*\*\**P* < 0.001).

### Pesticide exposure effects persist over time

In both species tested, the effect of treatment with either propoxur or chlorpyrifos was persistent over time. We tested survival following exposure to −16°C at 5-, 28-, and 55-days following dosing with either a solvent, propoxur, or chlorpyrifos. In solvent-treated ticks, there was no difference in survival over time for either species (Figure 2; *D. variabilis*: GLM: *F*_1,15_=0.7782, *p*=0.394, *R. sanguineus*: GLM: *F*_1,14_=0.06297, *p*=0.8058). *Rhipicephalus* ticks had a significant decrease over time in survival when treated with propoxur (Figure 2; GLM: *F*_1,13_=26.48, *p*=0.000188) but not chlorpyrifos (Figure 2; GLM: *F*_1,13_=0.8187, *p*=0.382). For *D. variabilis*, there was a significant decrease in survivorship in ticks treated with chlorpyrifos (Figure 2; GLM: *F*_1,13_=8,084, *p*=0.0138), primarily driven by extremely high survival (nearly 100%) 5 days post-exposure. There was a significant decrease in survival over the 55 day period in *D. variabilis* exposed to propoxur (Figure 2; GLM: *F*_1,13_=10.64, *p*=0.00618). Survival in pesticide-treated groups was significantly higher overall than in those treated with solvents over time (Figure 2; GLM: *F*_5,26_=2.462, overall, *p*=0.00723).

**Figure 2:**
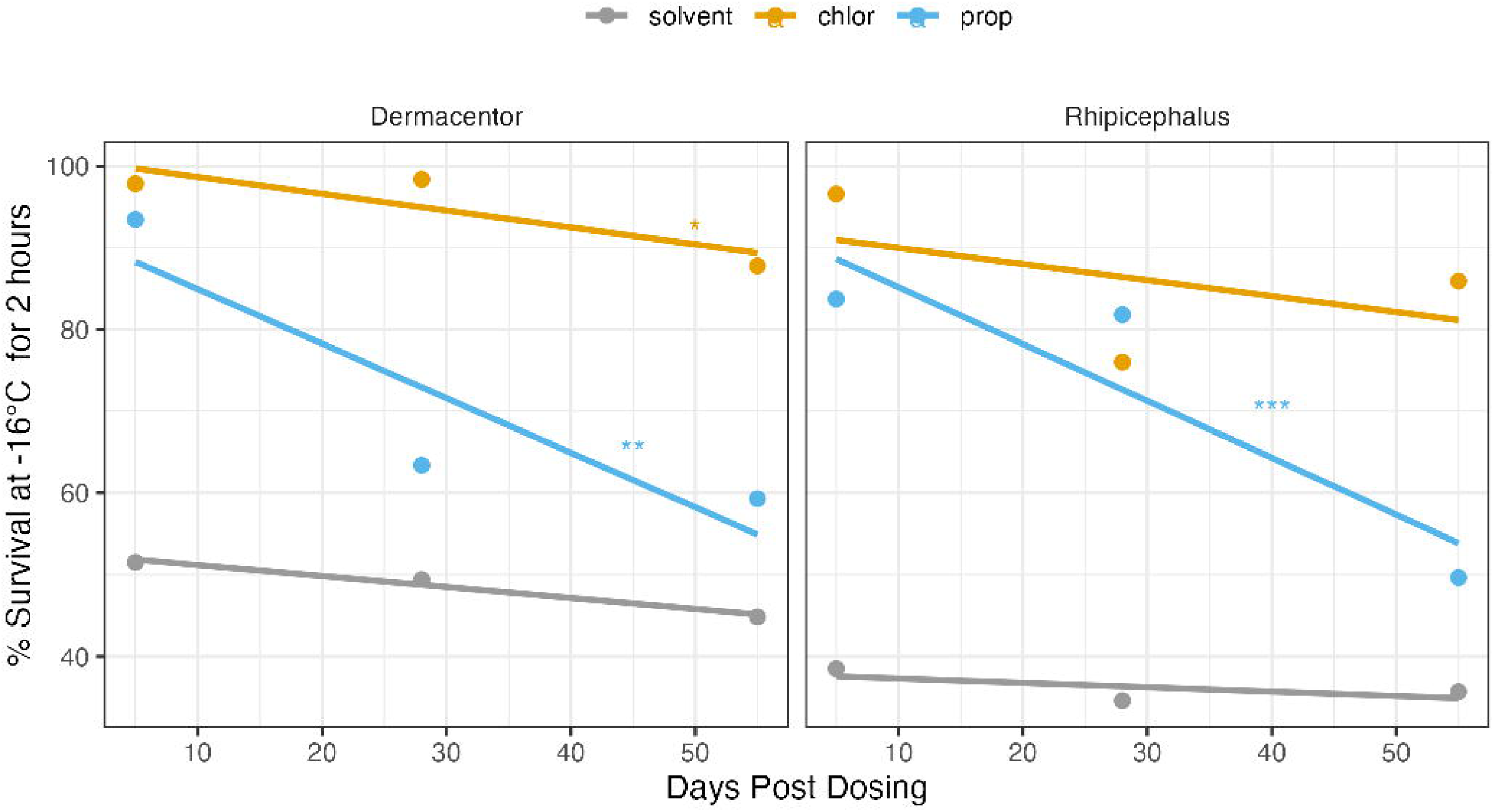
The proportion of *Dermacentor* (A) and *Rhipicephalus* (B) that survived a 2-hour exposure to −16°C at 5-, 28-, and 55-days post treatment with either a solvent (grey), propoxur (blue), or chlorpyrifos (yellow). Data points represent the mean survival of 5 repeated trials with 20 individuals per trail. Lines are linear models in the form of: survival ∼ days_post_treatment, where days_post_treatment is a continuous variable. Symbols indicate lines with significant slopes (\**P* < 0.05, \*\**P* < 0.01, \*\*\**P* < 0.001), showing that survival significantly decreased over time since exposure.

### Establishing LT_50_ for pesticide-treated D. variabilis

Survival declined significantly with colder temperature exposures in all ticks (Figure 3; GLM: *F*_1,72_=111, overall, *p*<0.00001), allowing the determination of lethal temperature estimates for each treatment group. Both the propoxur- and chlorpyrifos-treated groups were significantly different from the solvent-treated group overall, having generally higher survival than controls (Figure 3; GLM: *F*_3,70_=62.57, propoxur *p*=0.0025; chlorpyrifos *p*<0.0001). Within each treatment temperature, chlorpyrifos and propoxur treated ticks had significantly higher survival than solvent-treated ticks except at −25°C because propoxur was not different from solvents and at - 28°C neither propoxur nor chlorpyrifos were significantly different from the solvent (Figure 3; chlorpyrifos: estimate = −4, *p*=0.5304; propoxur: estimate = 0, *p*=1.000). Solvent-treated ticks were the most susceptible to cold temperatures (LT50 = −15.68°C), followed by propoxur treated ticks (LT50 = −17.92°C), and chlorpyrifos (LT50 = −19.35°C).

**Figure 3:**
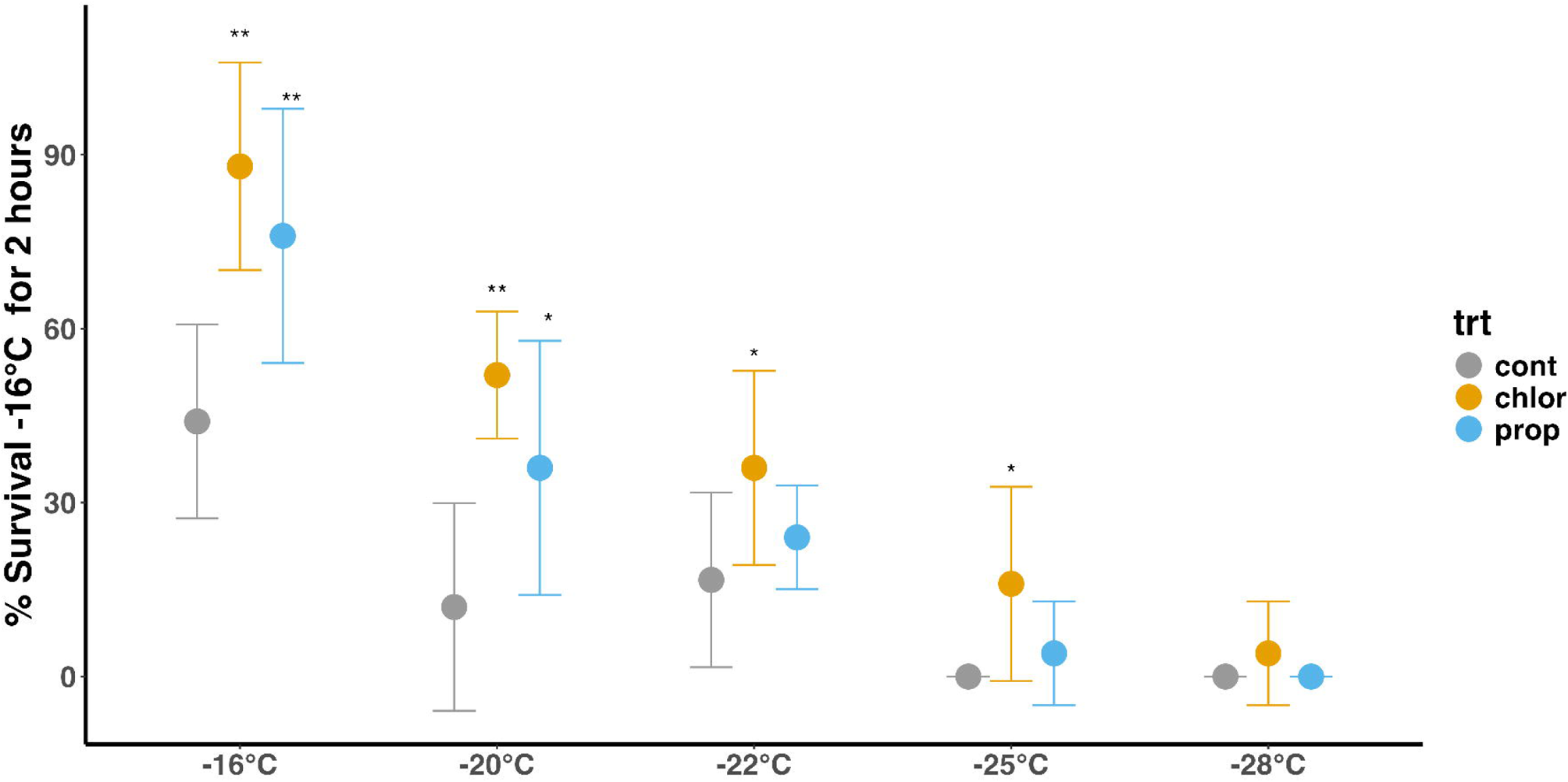
The percent survival of *D. variabilis* treated with either a solvent (grey) chlorpyrifos (yellow) or propoxur (blue) after a 2-hour low temperature exposure. Solvent-treated ticks were most susceptible to low temperatures (LT50 = −16°C). For ticks treated with chlorpyrifos and propoxur LT50s were significantly lower at −18°C and −19°C respectively. See text for statistical details. Data points show the mean survival of 5 repeated trials with 5 individuals per trail with standard errors.Asterisks indicate values that were significantly different from solvent controls (t-test, Tukey method, \**P* < 0.05, \*\**P* < 0.01, \*\*\**P* < 0.001).

### Pesticide exposure increases overwintering survival at northern sites for D. variabilis

At the southern site in Cincinnati OH, USA, overwintering survival was significantly greater in the chlorpyrifos-treated ticks than the solvent-treated ticks, but there was no significant difference between the propoxur- and solvent-treated ticks (Figure 4; chlorpyrifos: estimate = - 0.1745, *p*=0.0203; propoxur: estimate = −0.0605, *p*=0.7987). At the northern site in Crooks, SD, USA, ticks treated with chlorpyrifos or propoxur before overwintering had significantly greater survival than those treated with a solvent (Figure 4; chlorpyrifos: estimate = −0.1345, *p*=0.0050; propoxur: estimate = −0.0405, *p*=0.0245). Treatment with propoxur and chlorpyrifos increased survival from approximately 37% in the control group to 43% and 60% respectively in the northern site. In general, survival was significantly higher among all treatment groups in the southern site, compared with the northern site, suggesting that overwintering temperatures may drive mortality in ticks at or near their northern range limits.

**Figure 4:**
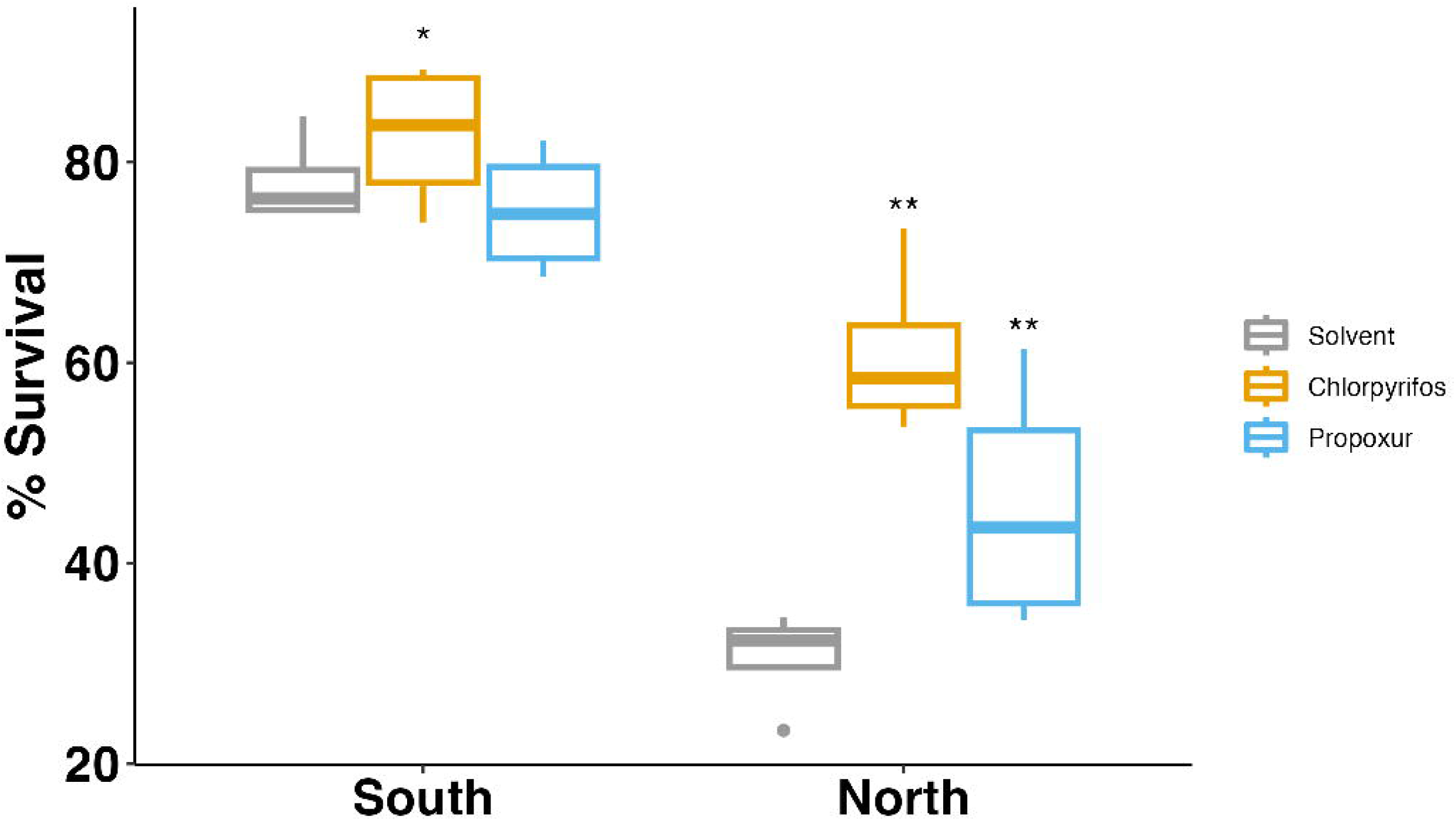
Overwintering survival varied significantly between ticks exposed to chlorpyrifos (yellow), propoxur (blue), and solvent (grey) in the northern site (Crooks, SD, USA), but not at the southern site (Cincinnati, OH, USA). Boxplots show mean survival in 4 sets of 100 ticks in each treatment group at each site. Asterisks indicate values that were significantly different from solvent controls for each site (t-test, Tukey method, \**P* < 0.05, \*\**P* < 0.01, \*\*\**P* < 0.001).

### Comparative RNA-seq analysis shows significant overlap between rapid cold hardening and pesticide exposure

When previous RNA-seq studies were evaluated for transcriptional differences, there was a significant overlap between expressional changes following rapid cold hardening and pesticide exposure (Figure 5). There were 1,793 contigs increased and 1,157 decreased following rapid cold hardening (RCH) (Figure 5a). Of those, 815 and 498 overlap with both pesticide exposures (chlorpyrifos and propoxur), which were increased and decreased, respectively. For those that increased, there was an enrichment for GO categories signaling, cytoskeleton reorganization, DNA integration, and intracellular transport (Figure 5b). There was a general suppression of the process of gene expression and translation associated with contigs that have reduced expression following rapid cold hardening and pesticide exposure (Figure 5b).

**Figure 5:**
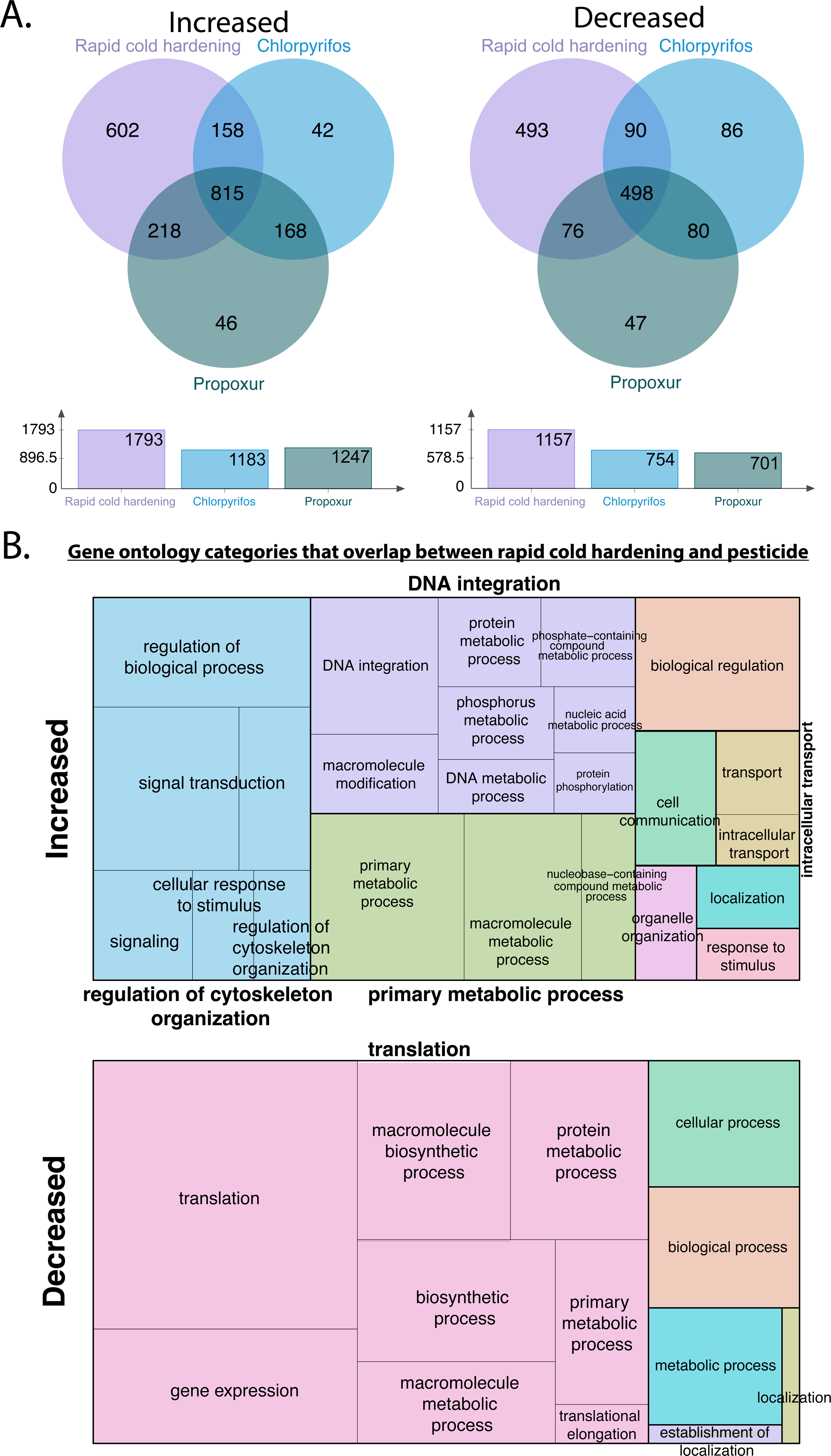
Impact of rapid cold hardening and pesticide exposure on gene transcription in American dog tick, *D. variabilis*. A. Three-way analysis to identify expression of genes that overlap for ticks exposed to cold and pesticides. B. Treemaps illustrating gene ontology (GO) terms for biological processes with overlapping increase following rapid cold hardening and pesticide exposure. Data from Pathak et al., 2022 and Rosendale et al., 2022.

### The impact of increased overwintering survival on current tick distributions

We compared the pSDMs which were constructed using the coefficients from the thermal response curves as priors, to the SDMs based on environmental data alone (Figure 6). Overall, the physiology SDMs had better fit than the correlative SDMs (pSDM: Boyce = 0.98; pesticide-pSDM: Boyce = 0.903; SDM: Boyce = −0.189; Table 1). This can primarily be attributed to the pSDMs showing more moderate occupancy in the north compared with the correlative SDMS (Figure 6). The correlative SDM had higher CCR (correct classification rate) and precision, but lower sensitivity, specificity and recall (sensitivity =0.833, specificity = 0.795, precision = 0.037, recall=0.833), compared with pSDMs (sensitivity =0.9, specificity = 0.764, precision = 0.034, recall=0.9) (Table 1) indicating that both model types accurately explained the measured distributions for *D. variabilis*. There was little difference in performance between pSDMs with and without the impacts of pesticides included (Table 1; Figure 6). Pesticide pSDMs showing current *D. variabilis* distributions had range limits extending slightly farther north compared with models not including pesticide impacts (Figure 6). Overall, the pSMDs outperformed correlative SDMs and showed a lower incidence of *D. variabilis* at northern latitudes.

**Figure 6.**
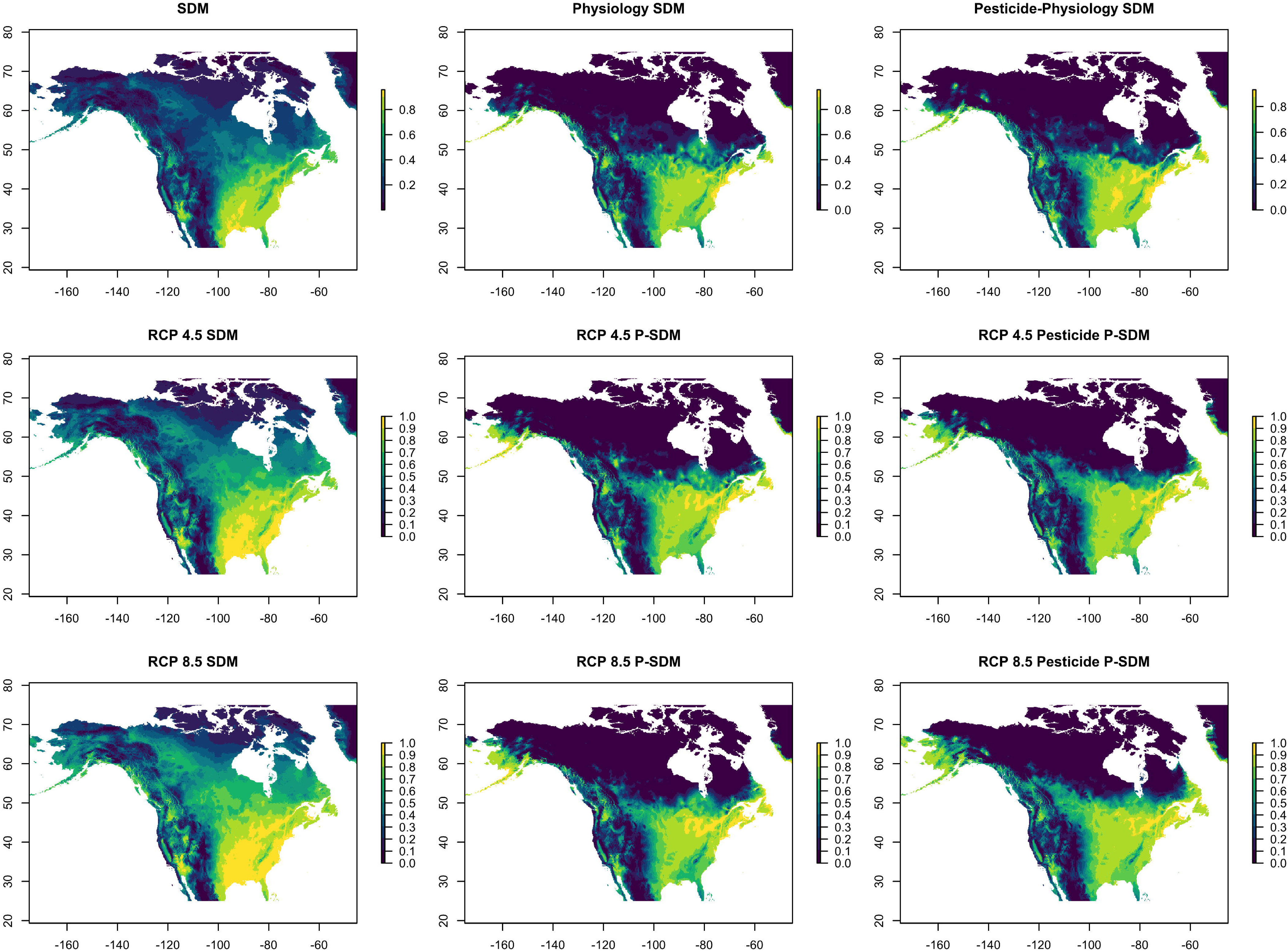
Predicted habitat suitability for *D. variabilis* in North America under current environmental conditions (top row) RCP 4.5 (middle row) and RCP 8.5 (bottom row). General linear models (row 1) were fitted to 425,390 occurrence data points for *D. variabilis*, or to interpolated probability estimates for occurrence given a predicted LT50 of −16°C for ticks unexposed to pesticides (column 2) or a predicted LT50 of −19°C for ticks exposed to pesticides (column 3).

**Table 1.**
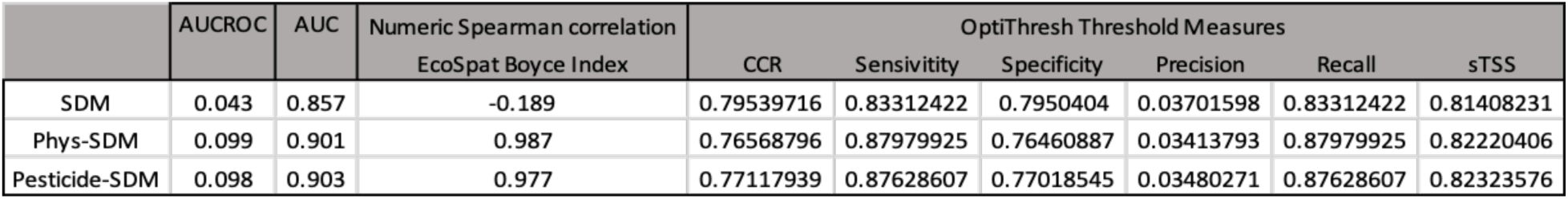
Model validation and performance estimates for correlative SDM, pSDM, and pSDM with pesticides.

### The impact of increased overwintering survival on future tick distributions

Under future climate scenarios (RCP4.5 and RCP8.5) *D. variabilis* will likely undergo a range shift to more northern environments (Figure 7). This was true for both correlative and physiology SDMs (Figure 7, Table 2). Both pSDMs and correlative SDMs had high overlap in RCP 4.5 (Figure 7, Table 2, for SDMS: Schoener D = 0.929, Warren I = 0.99, Hellinger D = 0.088; for pSDMs: Schoener D = 0.830, Warren I = 0.970, Hellinger D = 0.243; for pesticide pSDMs: Schoener D = 0.833, Warren I = 0.964, Hellinger D = 0.267). In RCP 8.5 the overlap was less, primarily due to greater northward range shifts (Figure 7, Table 2, for SDMs: Schoener D = 0.883, Warren I = 0.99, Hellinger D = 0.142; for pSDMs: Schoener D = 0.751, Warren I = 0.938, Hellinger D = 0.353; for pesticide pSDMs: Schoener D = 0.735, Warren I = 0.919, Hellinger D = 0.403). The overlap for pSDMs was less than the overlap for correlative SDMs which is most likely due to the correlative SDMs overestimating the current presence of *D. variabilis* in the north (Figure 7).

**Figure 7:**
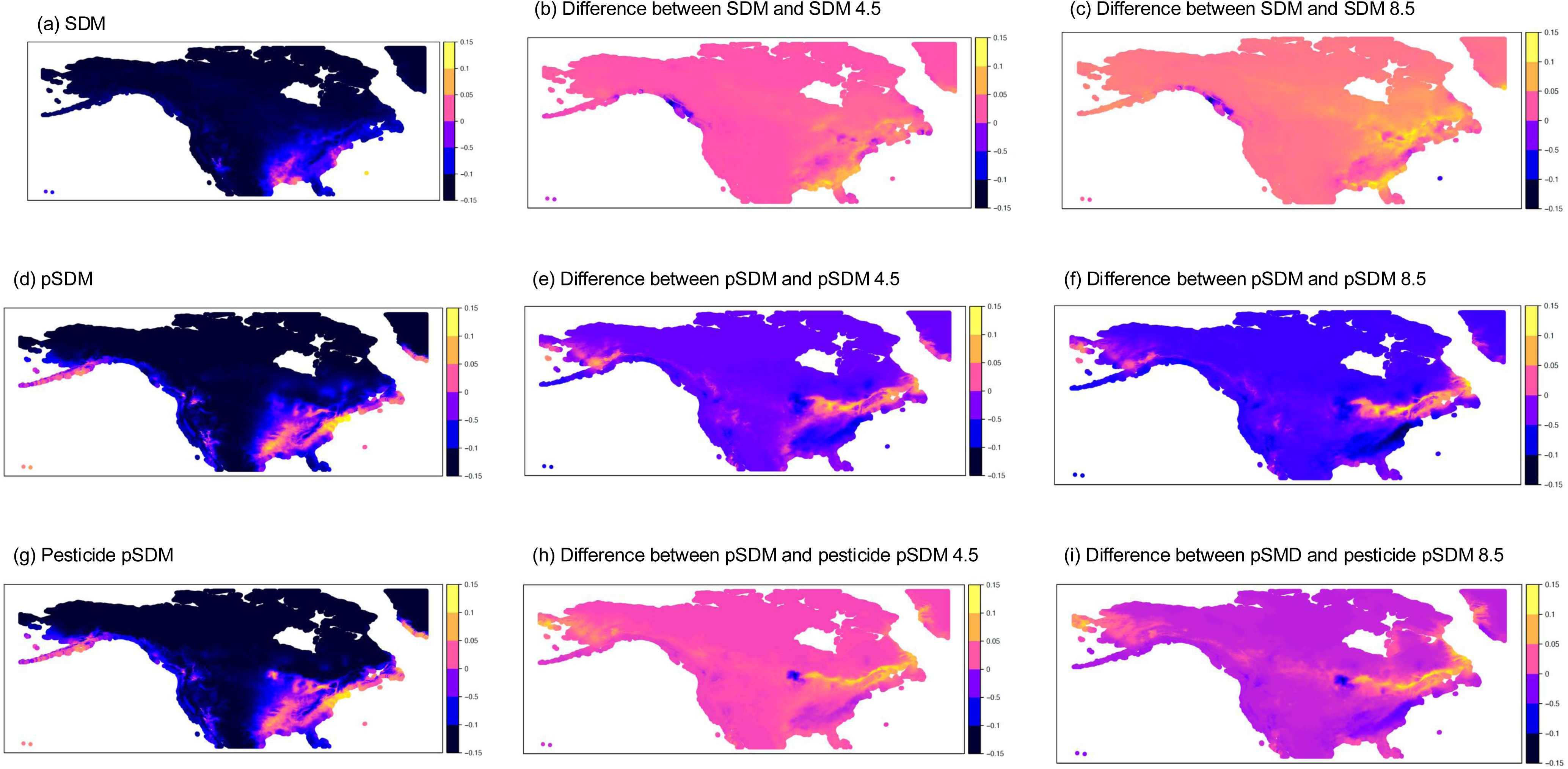
A-I represents the difference between the current (A, D, G) and future (B, E, H, C, F, I) distributions for *D. variabilis*. Blue colors indicate where models predicted lower suitability and yellow indicate regions where models predicted higher suitability. First column shows current predicted distributions for climate (a) physiology (d) and pesticide-physiology (g) models. The second column shows differences between current and future (RCP 4.5) distributions and the third column shows differences between current and future (RCP 8.5) distributions. Warm colors indicate regions where physiology-based models predict higher suitability than correlative SDMs.

**Table 2:**
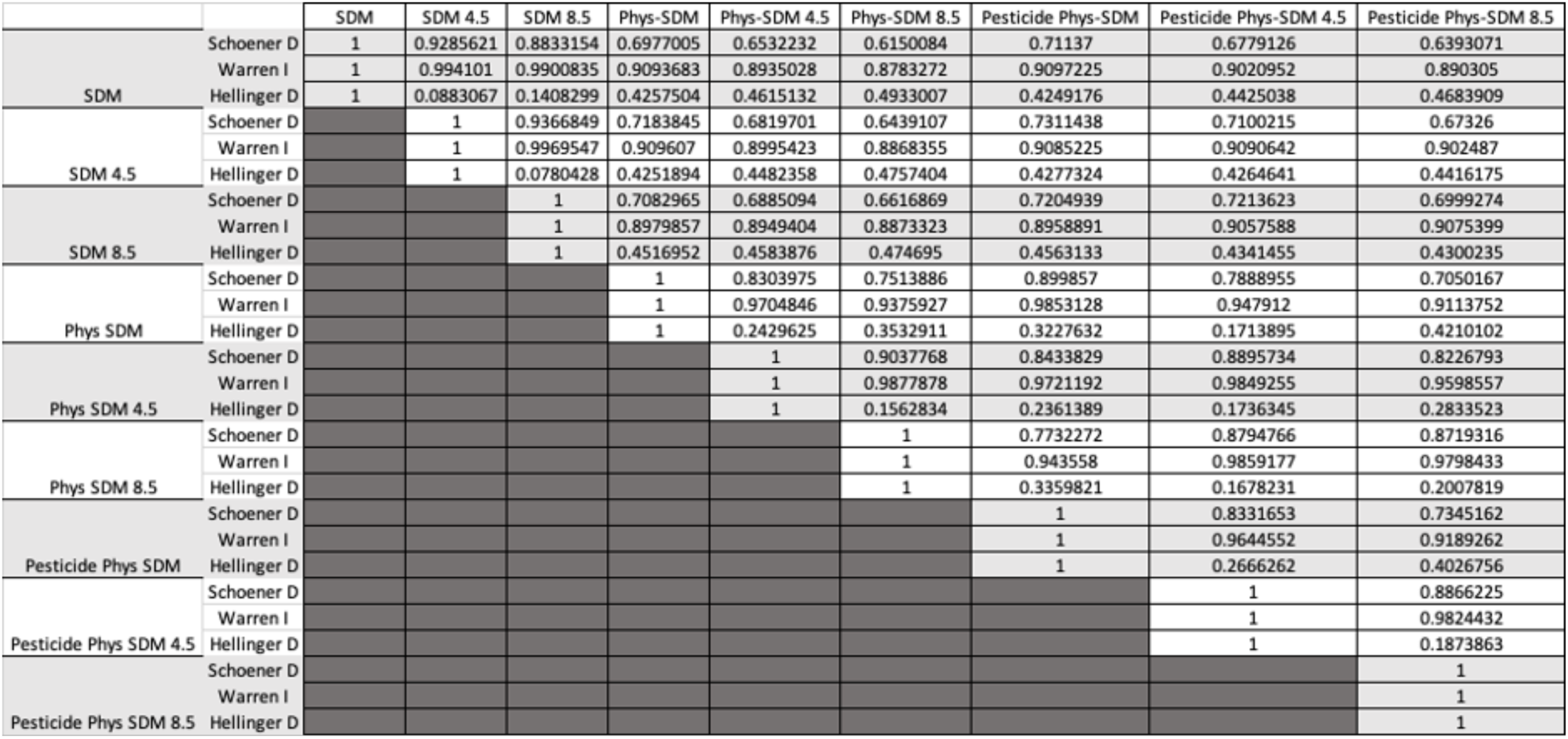
Overlap indices between pSDMs and correlative SDMs.

## Discussion

Our study combined lab, field, and theoretical approaches to measure the impact of pesticides on cold tolerance, overwintering survival, and distribution of a common pest. The results suggest that sublethal pesticide exposure alters tick cold tolerance, leading to higher overwintering survival and probably increases survival at northern range limits. RNA-seq studies between the process of RCH and pesticide exposure revealed a suite of factors underlying cross-tolerance to these stresses. Although several studies have shown that multiple stressors can lead to additive or antagonistic effects and unpredictable outcomes, less is known about the combined impact of pesticides and extreme temperatures. Environmental change and increased pesticide usage will lead to more significant overlap of these stressors for both target and non-target species, increasing the urgency of understanding potential interactions.

We demonstrated that the combined effects of pesticides and cold temperatures can increase overwintering survival, and permit greater northward range shifts by incorporating physiological data into SDMs. Although both correlative and physiological models correctly predicted current species’ ranges, the mechanistic models moderately outperformed the purely correlative models. The better fit of the physiological models was primarily driven by differences at the species’ range limits where physiological models tended to more precisely estimate the edges of current distributions. The increased survival at the southern site compared with the northern site suggests that cold temperatures are likely limiting northern populations of ticks. This is mirrored by the better performance of the physiological SMDs at the northern range limit. Additionally, physiological models tended to moderate the estimated impact of temperature change, which has been previously reported in other studies comparing correlative and physiological modeling approaches (Gamliel et al., 2020). Physiological species distribution models may therefore provide a valuable tool for accurately estimating the impact of environmental change on species’ ranges and particularly for pests and vectors which are likely to have significant effects on host populations as they invade new areas.

Previous studies have used species thermal performance traits to estimate responses to temperature variation, but these traits are likely impacted by several factors, including local adaptation and plasticity (Ayrinhac et al., 2004; Gunderson et al., 2017). Here, we found that cold tolerance in ticks is impacted by previous exposure to sublethal pesticides. Pesticides have systemic impacts on several biochemical pathways that influence cold tolerance, including the electron transport chain, ion regulatory channels, and cellular pumping (Stenersen, 2004). Maintenance of ion homeostasis across cellular space is critical for preserving muscle function at cool temperatures (MacMillan et al., 2015; MacMillan & Sinclair, 2011). Disruptions in ion regulatory channels leads to systemic depolarization across neurotransmitters and probably underlies the critical loss of muscle function at low temperatures (Robertson et al., 2017; Spong et al., 2016). Organophosphates and carbamates, which were tested in this study, operate on highly conserved biochemical pathways, including the release of neuroregulatory chemicals such as acetylcholinesterase. In the absence of a pesticide, acetylcholinesterase binds to acetylcholine and limits signaling at the synaptic cleft, but during exposure to organophosphates or carbamates, acetylcholinesterase is inhibited, leading to hyperactivation of the receptors (Aktar et al., 2009). Contrastingly, cold temperatures dampen signaling of neurotransmitters by slowing down the rate of molecular pumps and movement at the synaptic cleft (Robertson & Money, 2012). We predict that the confounding effects of dampening at cold temperatures and hyperactivation of pesticides have an antagonistic effect, leading to better muscular performance in the cold. More work is needed to vet this prediction and determine the molecular mechanisms underlying these interactions.

Another non-exclusive explanation for the observed increase in cold tolerance after pesticide exposure is that pesticide exposure may trigger the production of stress proteins, which also provide protection from extreme temperatures. This effect has been observed in mosquitoes (Mack & Attardo, 2024). We tested this possibility by comparing the differential gene expression of ticks exposed to cold (Rosendale et al., 2022) and pesticides (Pathak et al., 2022). For ticks to survive environmental stressors associated with winter, such as low temperatures and dehydration, processes such as rapid cold hardening are likely to be critical to increased survival, which has been observed in the American dog tick (Rosendale et al., 2016). Cross-tolerance between stress types is commonly observed in arthropods (Boardman, 2024; Kaunisto et al., 2016), and has been observed between dehydration and cold exposure for ticks (Rosendale et al., 2016). Pesticide exposure has been shown to cause drastic transcriptional shifts in ticks (Pathak et al., 2022), suggesting potential overlap in gene regulation between pesticide exposure and rapid cold hardening. Substantial overlapping molecular pathways were observed based on previously available RCH and pesticide exposure RNA-seq (Rosendale et al. 2022; Pathak et al. 2022). Specifically, RCH and pesticide exposure in ticks involve the activation of stress-responsive mechanisms, such as signal transduction, DNA integration, cytoskeletal regulation, intracellular transport, and the suppression of protein translation (Pathak et al., 2022; Rosendale et al., 2022). These factors are likely critical to the improved cold tolerance following pesticide exposure. As examples, DNA repair mechanisms are crucial in cold and pesticide stress as DNA damage is typical, and DNA integration is vital to ensure repair (Hernández-Toledano & Vega, 2023; Lubawy et al., 2019). Cytoskeleton shifts play a key role in maintaining cell structure and function under stress, which have been identified as associated with overwintering and cold stress (Kim et al., 2006; van Oirschot & Toxopeus, 2025; Wang et al., 2023). Lastly, translation is suppressed in both processes, suggesting that reduction in this machinery underlying protein synthesis may be critical to allowing time for recovery before resuming normal biological processes (Wang et al., 2023). These shared responses suggest that ticks utilize overlapping transcriptional shifts to respond to cold and pesticide exposure, which are likely critical to the increased cold tolerance and overwintering survival of ticks after pesticide exposure that we observed.

Predicting the impact of temperature on organism survival will likely depend on close measurement of several interacting stressors, which include natural environmental factors (thermal and dehydration) in conjunction with those that are artificial (pesticides and pollutants). The interaction of multiple stressors will likely drive species’ responses to environmental change, and incorporating these data into physiological models will likely improve predictions. Mechanistic models consistently outperformed correlative models in our study. This suggests that understanding the species-specific physiology is critical to accurately predicting future pest distributions and provides a valuable tool for allocating resources towards the most vulnerable human and animal populations.

## Supporting information

Supplemental Figure 1: Mean daily temperatures at overwintering exclosure sites for Cincinnati OH, USA (pink) and Crooks, SD, USA (blue).

Supplemental Figure 2: Data used for physiological sub-models. The proportion of adult D. variabilis alive following 2-hour temperature exposures for

Supplemental Figure 2: Data used for physiological sub-models. The proportion of adult D. variabilis alive following 2-hour temperature exposures for

## Funding and Acknowledgments

KJO was partially supported by the David H. Smith Postdoctoral Fellowship and by the United States Department of Agriculture, Agricultural Research Service (#2090–32000-040-000D). Research reported in this publication was supported partially (reusable equipment) by the National Institute of Allergy and Infectious Diseases under Award Numbers R01AI148551, R21AI166633, and R21AI176098 (JBB). Mention of trade names or commercial products in this publication is solely to provide specific information and does not imply recommendation or endorsement by the USDA. The USDA is an equal opportunity provider and employer.

## Declarations

### Consent for publication

All authors have consented for the publication of this manuscript. Availability of data and materials: The datasets used and/or analyzed during the current study are available from the corresponding author on reasonable request.

## SUPPLEMENT

**Supplemental Figure 1:** Mean daily temperatures at overwintering exclosure sites for Cincinnati OH, USA (pink) and Crooks, SD, USA (blue).

**Supplemental Figure 2:** Data used for physiological sub-models. The proportion of adult *D. variabilis* alive following 2-hour temperature exposures for control group (A) and chlorpyrifos-treated group (B) with logistic regression lines. The logistic regression model has the form:

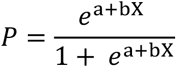

Where P is the probability of an event (e.g., the probability of survival) and e is the basis of the natural logarithm, and a and b are the parameters of the model (in this case the slope and regression of the best fit line for the survival data).

**Supplemental Figure 3:** The proportion of alive adult *D. variabilis* following exposure to - 16°C among untreated, water-treated, and solvent-treated ticks. Boxplots represent 7 sets of 20 individuals measured in parallel with each experiment described in methods sections.

