## Supplementary figures and images for "Sub-lethal pesticide exposure facilitates the potential northward range shifts of ticks by increasing cold tolerance and overwintering survival"

### Supplemental Data 1

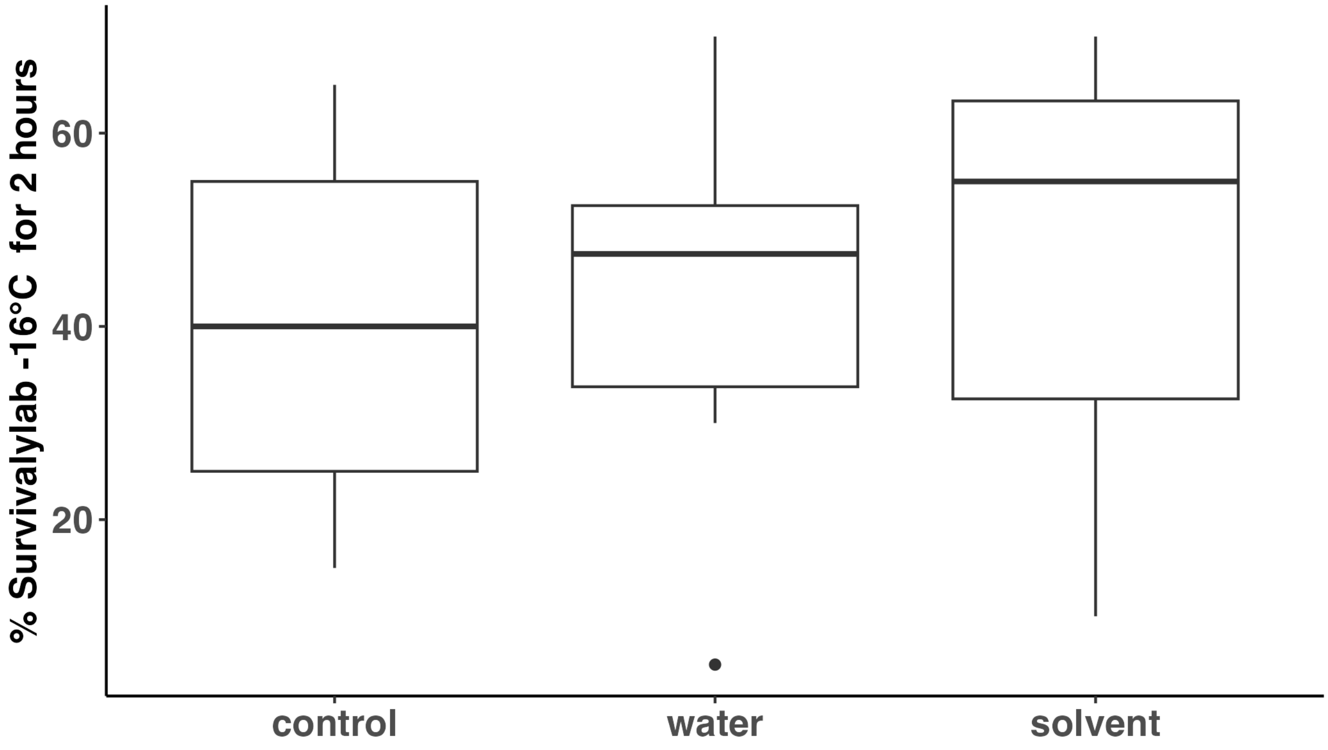

### Supplemental Figure 1: Mean daily temperatures at overwintering exclosure sites for Cincinnati OH, USA (pink) and Crooks, SD, USA (blue).

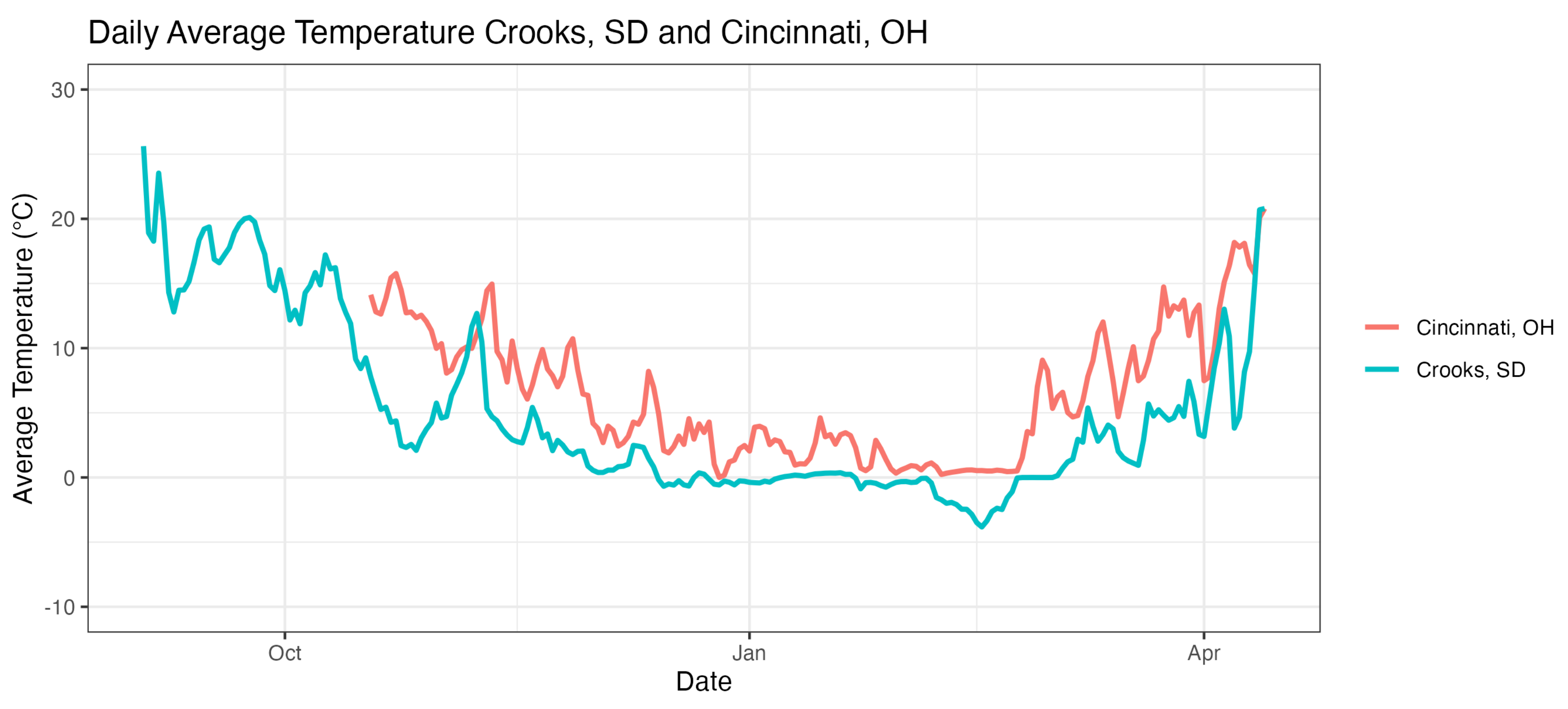

### Supplemental Figure 2: Data used for physiological sub-models. The proportion of adult D. variabilis alive following 2-hour temperature exposures for

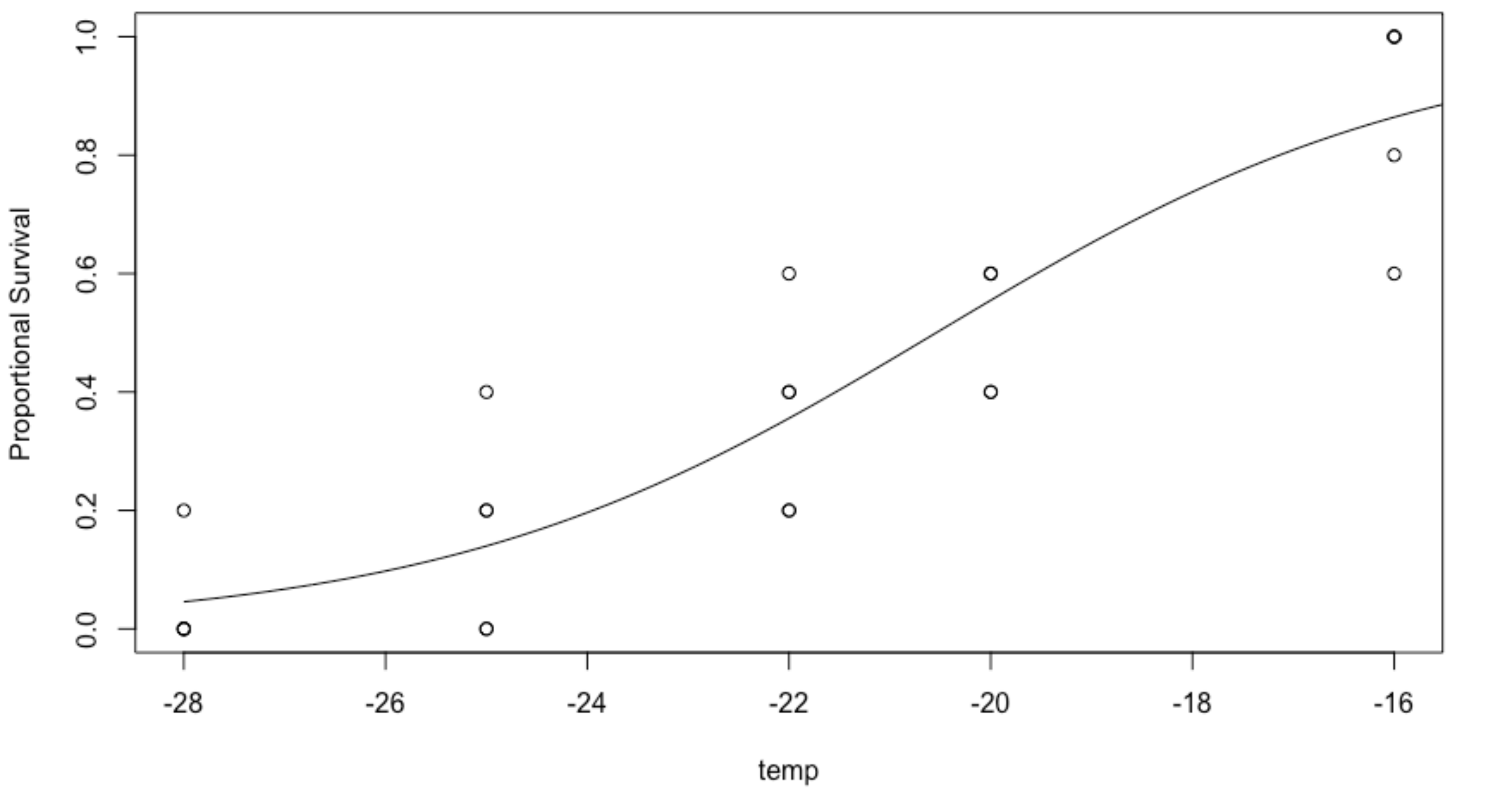

### Supplemental Figure 2: Data used for physiological sub-models. The proportion of adult D. variabilis alive following 2-hour temperature exposures for

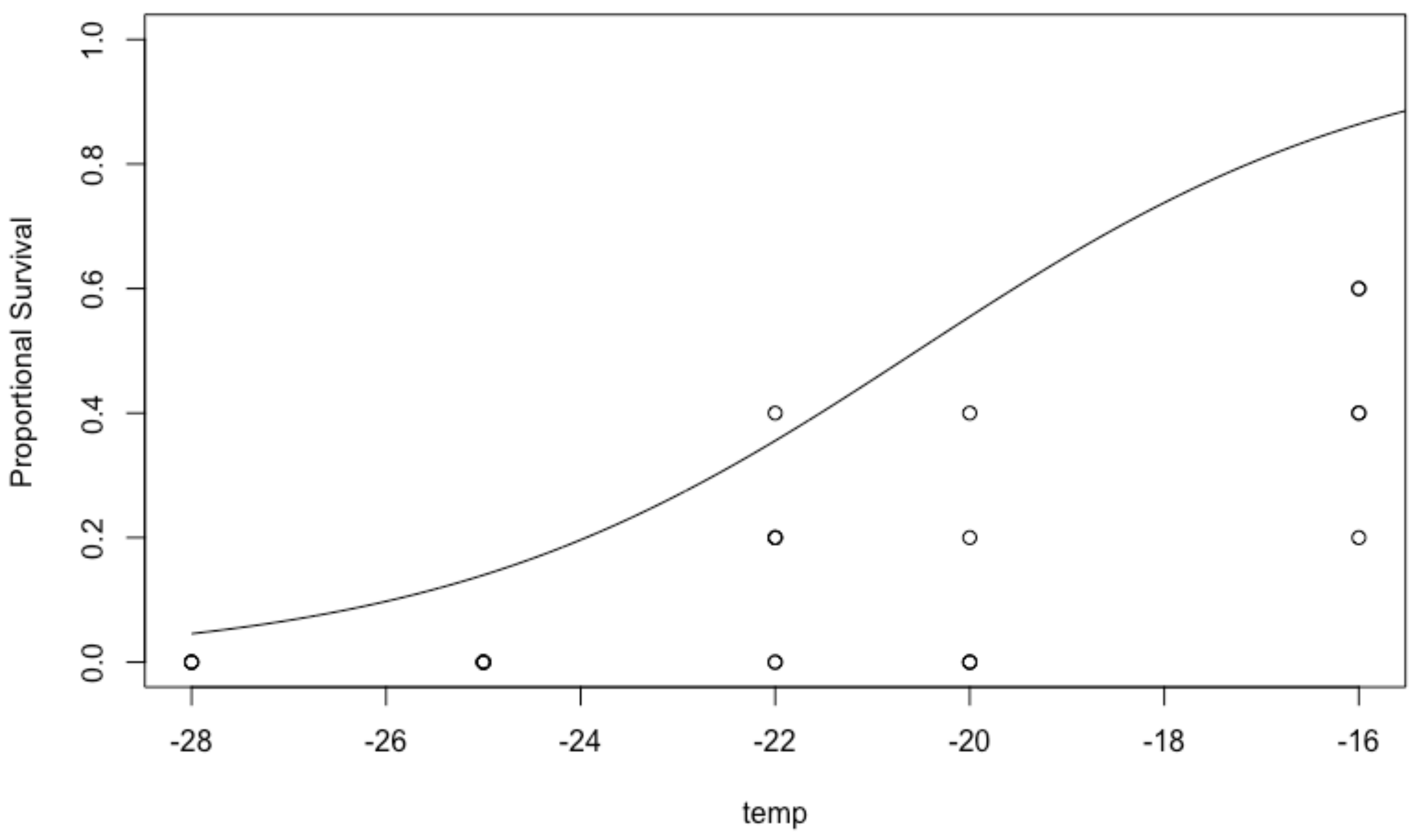
